# The development of FEDUPP: Feeding Experimentation Device Users Processing Package to Assess Learning and Cognitive Flexibility

**DOI:** 10.1101/2025.08.14.670424

**Authors:** Mingyang Yao, Avraham M. Libster, Shane Desfor, Freiya Malhotra, Nathalia Castorena, Patricia Montilla-Perez, Francesca Telese

**Author notes:** equal contribution (Mingyang Yao, Avraham M. Libster).

## Abstract

Cognitive flexibility, the ability to adapt behavior in response to changing contingencies, is a key component of adaptive decision-making and is impaired in multiple neuropsychiatric disorders. Traditional rodent assays of cognitive flexibility are conducted in experimenter-controlled sessions in restrictive environments, limiting ecological validity and temporal resolution. Here, we developed a fully automated, home-cage paradigm using the Feeding Experimentation Device 3 (FED3) and a companion open-source analysis pipeline, the Feeding Experimentation Device Users Processing Package (FEDUPP), to assess learning and cognitive flexibility with minimal experimenter intervention. The paradigm combines a single-day fixed-ratio 1 (FR1) task with a multi-day, reversal learning task in which active port assignment switches every 25 pellets collected. FEDUPP implements multi-scale learning metrics, including overall accuracy, an 80% accuracy milestone, and a machine learning–based classification of meal accuracy to capture motivated, goal-directed feeding. In wild-type mice, the paradigm detected rapid FR1 acquisition and progressive within-block adaptation during reversal. Application to mice with dorsal hippocampal knockdown of the scaffolding protein CASK revealed faster FR1 acquisition and higher accuracy than controls but a delayed onset of the first accurate meal after reversal, suggesting a selective deficit in updating goal-directed feeding behavior. These findings demonstrate that FEDUPP enables high-resolution, continuous assessment of learning and cognitive flexibility in ethologically relevant settings, and that meal-based accuracy provides a sensitive metric for detecting subtle flexibility impairments not captured by traditional measures.

## Introduction

Cognitive flexibility is the ability to adapt behaviors and thought processes in response to changing environments, enabling individuals to revise strategies, switch tasks, and adopt new response patterns when reward structures or task demands shift (Dajani & Uddin, 2015; Diamond, 2013; Egner, 2023). This adaptability is necessary when suppressing previously learned behaviors to achieve new goals (Bari & Robbins, 2013).

Underlying cognitive flexibility is a network of neural processes that enable the efficient and dynamic allocation of cognitive resources (Shenhav et al., 2013). While much of the focus has traditionally been on the prefrontal cortex and basal ganglia (Cools, 2016; Dalley et al., 2004; Kim et al., 2011; Shenhav et al., 2013), the hippocampus also plays a significant role due to its involvement in contextual processing (Eichenbaum, 2017). Deficiencies in cognitive flexibility contribute to maladaptive behaviors seen in disorders such as schizophrenia, obsessive-compulsive disorder, and addiction, highlighting the importance of understanding its underlying neural mechanisms (Gruner & Pittenger, 2017; Izquierdo & Jentsch, 2012; Waltz, 2017).

Numerous behavioral assays have been developed to investigate the neural basis of cognitive flexibility in both human and animal models. These methods are based on reversal learning paradigms and involve adapting behavior when previously rewarded cues are reversed (Izquierdo et al., 2017; Robbins, 1481). These include the Wisconsin Card Sorting Test (WCST) (Bissonette et al., 2013; Milner, 1963), T-maze and Y-maze tasks (Dorofeikova et al., 2021; Ragozzino, 2007), instrumental discrimination reversal tasks (Birrell & Brown, 2000; Klanker et al., 2013; Rogers et al., 2000), and operant chamber tasks in contexts of probabilistic reward shifts (Floresco et al., 2008, 2009). These methods have demonstrated the important roles of the prefrontal cortex and nucleus accumbens in regulating flexible behavior (Dalton et al., 2016; Klanker et al., 2013). Despite the valuable insights gained from these methods, they come with limitations.

Behavioral assays have been conducted within experimenter-controlled sessions in restrictive environments, which can affect natural learning and behavior due to factors such as handling-related stress and the novelty of unfamiliar environments. Moreover, traditional behavioral analyses in cognitive flexibility research often rely on summary metrics such as cumulative accuracy or error rates, which do not fully capture the dynamics of learning and adaptation over time (Dajani & Uddin, 2015). These metrics often fail to represent the continuous process of behavioral adjustment and may overlook critical transitions between different states, such as exploration, where new strategies are tested to gather information, and exploitation, where known successful behaviors are used to maximize rewards (Diamond, 2013). Finally, manual video scoring methods introduce potential inaccuracies due to human error and subjectivity, limiting the resolution of complex behaviors (Anderson & Perona, 2014). This limitation underscores the need for more sophisticated methods to analyze nuanced behavioral adaptations, which can reflect complex cognitive processes like reinforcement learning involving trial-and-error learning to optimize actions and flexible decision-making that balances exploration and exploitation.

To address these challenges, the Feeding Experimentation Device 3 (FED3) was utilized in a home-cage environment, allowing for continuous, minimally invasive tracking of operant learning and behavioral adaptation (Matikainen-Ankney et al., 2021; Reichenbach et al., 2022). In addition, a behavioral assay protocol and accompanying software toolbox, FEDUPP (Feeding Experimentation Device Users Processing Package) was developed, to assess learning and cognitive flexibility. This allowed animals to autonomously engage with reversal tasks in their home cage, preserving naturalistic behaviors across circadian cycles and reducing stress-related confounds. Furthermore, new metrics to evaluate learning efficiency and transitions were established. To that end, machine learning for detailed behavior classification was used.

Following the establishment of the assay and analysis framework, they were validated under conditions of known neural disruption. For this, male C57BL/6 mice underwent targeted knockdown (KD) of CASK (Calcium/Calmodulin-Dependent Serine Protein Kinase) in the dorsal hippocampus via shRNA-mediated gene silencing (CASK-KD). Littermate controls received hippocampal injections of a non-targeting shRNA construct (Control). This perturbation model enabled the assessment of the sensitivity of behavioral assays and analytical metrics in detecting cognitive flexibility deficits resulting from molecular disruption.

These innovations provide a refined and scalable approach for investigating cognitive flexibility and adaptive learning in animal models, capturing the complexity of learning processes crucial for informed decision-making.

## Results

### Development of a FED3-based behavioral paradigm and analysis pipeline to assess cognitive flexibility

To assess cognitive flexibility in a home cage environment, FED3 devices (Matikainen-Ankney et al., 2021) were used to monitor mouse food intake over a two-phase behavioral paradigm. In the first phase, mice performed a nose-poke-based two-alternative forced-choice task with a fixed ratio of one nose poke per reward (FR1) for approximately 24 hours. In the second phase, mice completed a reversal version of the same two-alternative forced-choice task for approximately 72 hours (**Fig. 1**), in which the ‘active’ port switched after every 25 pellets collected (**Fig. 1**). The interval between port switches, defined by the time taken to collect 25 pellets, was termed a block. An in-house analysis pipeline, FEDUPP, was developed to extract learning and cognitive flexibility metrics from FED3 data. FEDUPP quantifies performance across multiple timescales, including overall accuracy, 80% accuracy milestone, and meal pattern. ‘Accuracy’ is defined as the proportion of correct (‘active’) port pokes relative to all port pokes (‘active’ + ‘inactive’), within a defined time window (FR1 session, behavioral block, meal) (**Fig. 2**, see methods). The ‘80% accuracy milestone’ was defined as the earliest time point at which accuracy exceeds 80% for a sustained 2-hour window, providing a shorter time scale measure of learning independent of the length of the entire session (**Fig. 2**, see Methods). In addition, sequences of ‘pellet collection’ events were identified as meals, and their measured accuracy was the meal’s accuracy (**Fig. 3** see methods). A long short-term memory (LSTM)-based classifier was trained on meals and their accuracy collections to classify meals into ‘accurate’ and ‘inaccurate’. The occurrence of the first ‘accurate’ meal was an additional evaluation of learning (**Fig. 3**, see Methods).

**Figure 1.**
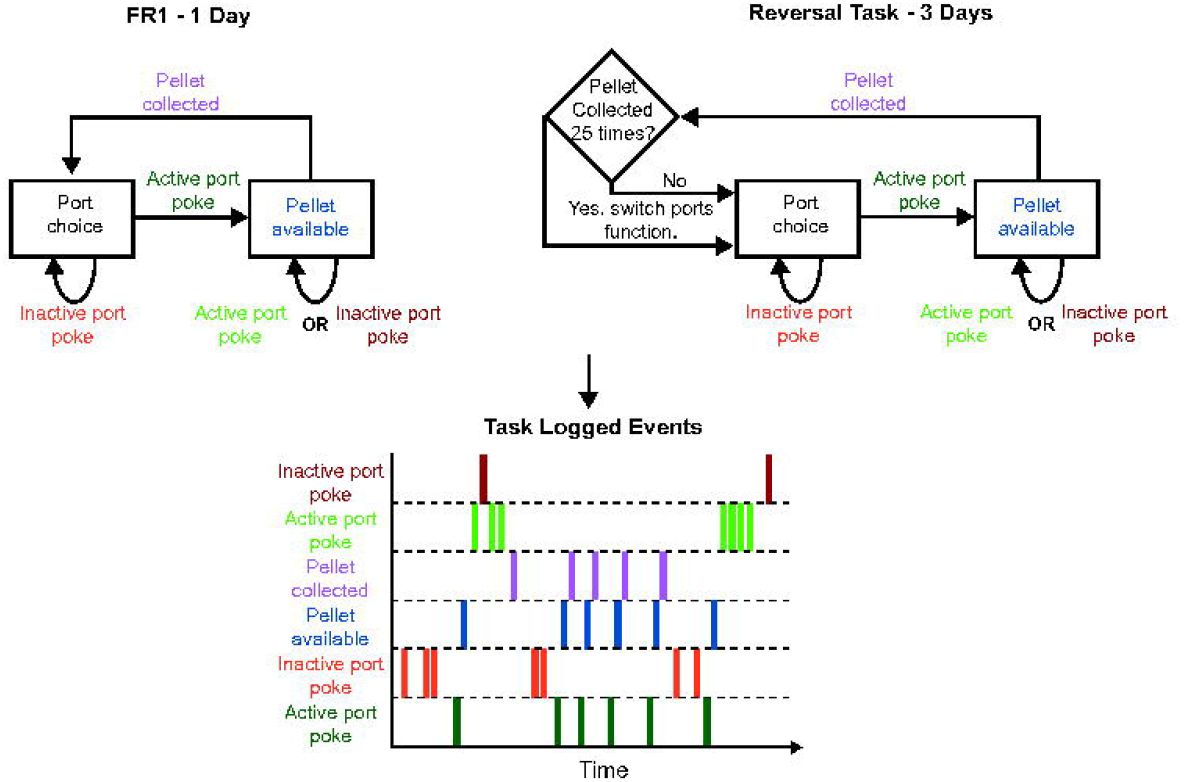
Schematic of FEDUPP Behavioral Assay and Analysis. State machine description of both the FR1 and Reversal tasks. Color-coded actions describe events logged by the FED3 device. On the bottom side, a schematic of the logged events is visualized as a time series.

**Figure 2.**
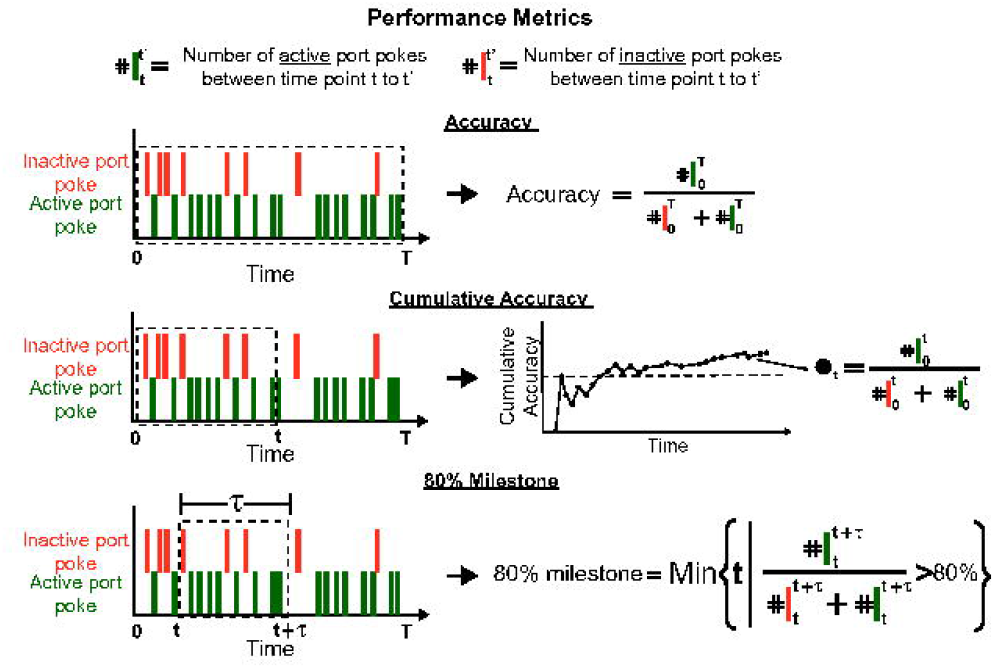
Schematic of FEDUPP calculations of performance metrics. upper panel - Given a time series of ‘active’ and ‘inactive’ port pokes (green and orange, respectively), the accuracy is calculated as ‘active’ divided by the sum of ‘active ‘and’ inactive’ port pokes events during the time period. Middle panel - Cumulative accuracy is calculated as the accuracy between 0 and a given time point t, where t is smaller than a given maximal value T. lower panel - 80% milestone is calculated as minimal value t, such that the accuracy in the time range t to t+tau (tau length of time window) is bigger than 80%.

**Figure 3.**
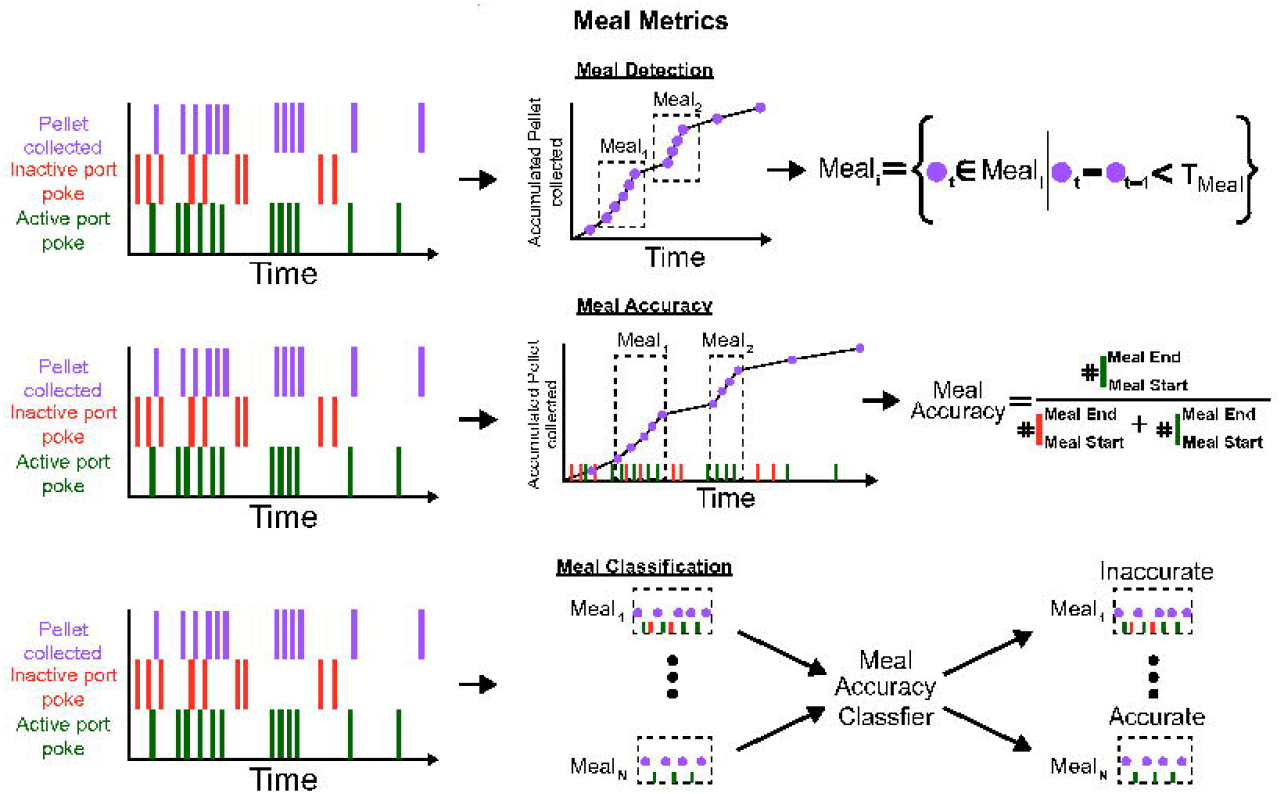
Schematic of FEDUPP calculations of meal metrics. upper panel - Given a time series of ‘active’, ‘inactive’ port pokes and ‘pellet collected’ events (green, orange and purple respectively), meal is defined as a series ‘pellet collected’ events in which the time difference between each on is smaller than a maximum defined by Tmeal. Middle panel - Meal accuracy is calculated as the accuracy based on the active and inactive port pokes events occurring during a meal. Lower panel - A meal is classified as either accurate or inaccurate based on the time series of the ‘pellet collected’ events and the meal’s accuracy.

A cohort of wild-type C57/Bl6 mice (WT) was used for initial method development. This group enabled the establishment of baseline performance metrics and refinement of computational analysis tools.

### Mice acquired above chance-level FR1 performance in less than 24 hours

In the FR1 session, WT mice collected an average of ∼209 pellets (209.30±8.30, N=17, **Fig. 4A**), corresponding to ∼4.2 g of chow. The mice improved their accuracy over time, reaching an average overall accuracy of ∼ 68% (68.28±1.8%, N=17, **Fig. 4B, C**) within the initial 24 hours. Notably, the 80% accuracy milestone was achieved after the first ∼5 hours (4.91±0.82 hours, N=17, **Fig. 4D**) from the start of the FR1. Moreover, using the pre-trained LSTM-based meal classifier (**Fig. 3**), the first “accurate” meal was detected at ∼ 2.7 hours (2.68±0.28, N=17, **Fig. 4E**).

**Figure 4.**
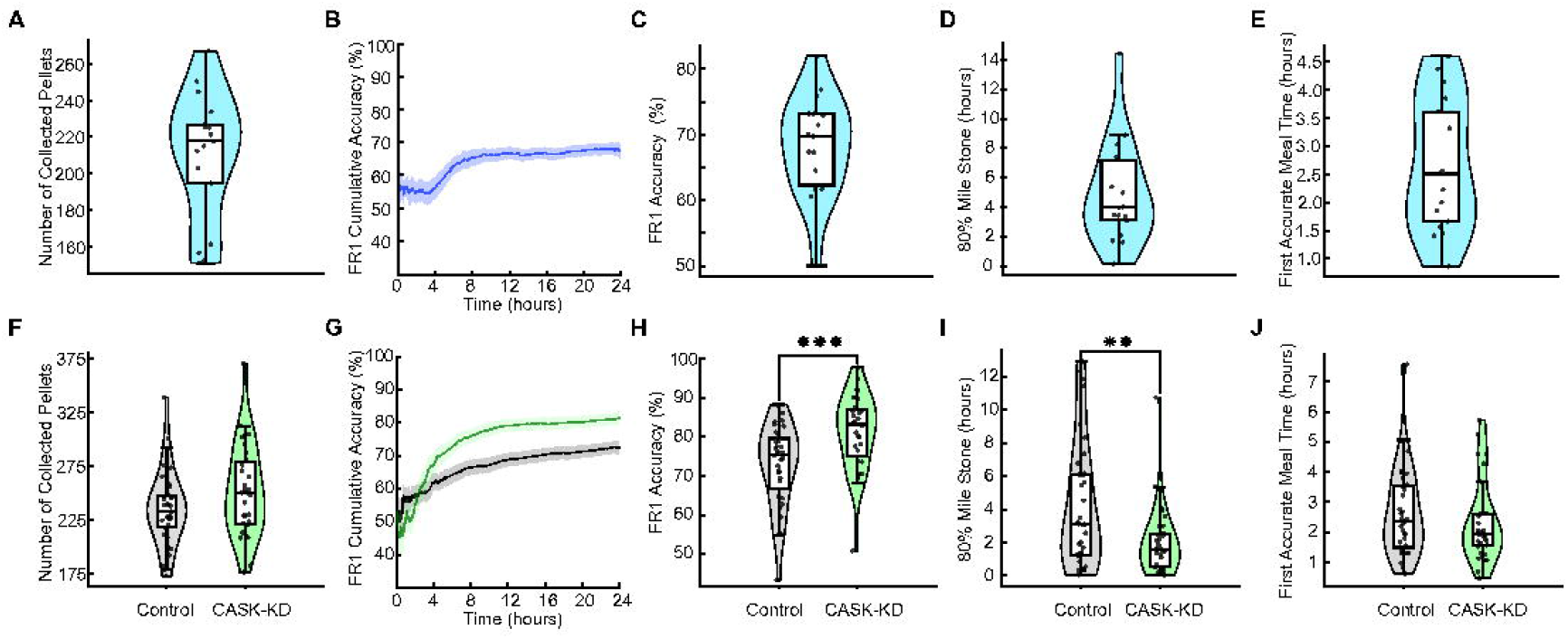
Behavioral Measures of the WT validation and CASK cohorts experiment during FR1 protocol. **A**,**F**. Number of collected pellets during the FR1 of the WT (A), Control and CASK-KD (F). **B**,**G**. Cohorts Dynamics of accuracy during the FR1 of the WT (B), Control and CASK-KD (G) cohorts during the 24 hours of the FR1 task. **C**,**H**. Total accuracy during the FR1 session for the WT (C) and CASK-KD (H). CASK-KD had a significantly higher accuracy than the Control. **D**,**I**. First time point to reach 80% accuracy in a time window of two hours of the WT (D), Control and CASK-KD (I) cohorts during FR1. **E**,**J**.First accurate meal time during the FR1 of the WT (E), Control and CASK-KD (J) cohorts. In panels F,H-J, Grey and Green designate Control and CASK-KD groups, respectively.

When applied to the CASK-KD and control groups, the FR1 paradigm revealed that the CASK-KD group showed a non-significant trend toward collecting more pellets than control group (CASK-KD: 251.3±7.48, N=31; Control: 235.46±5.76, N=35, p=0.100, **Fig. 4F**). Compared to control mice, the CASK-KD mice exhibited significantly higher overall accuracy (81.25%± 1.72 vs 72.74% ± 1.76, CASK-KD N=31, Control N=35, p = 0.001, **Fig. 4 G, H**) and required significantly less time to reach the 80% milestone(1.99±0.37 hours vs 4.22±0.63 hours, CASK-KD N=31, Control N=35 (p = 0.005, **Fig. 4I**). In contrast, a comparison of the first accurate meal between the CASK-KD and control groups revealed no differences between groups (CASK-KD: 2.31±0.23 hours, N=31; Control: 2.63±0.24 hours; N=35, p = 0.172, **Fig. 4J**).

Across all cohorts, performance exceeded chance level by the end of the FR1 phase, with CASK-KD mice showing an increased accuracy and faster acquisition relative to controls, as reflected in the 80% milestone metric.

### Mice perform a reversal paradigm with above-chance accuracy during each block

Following the FR1 assay, mice completed a reversal task lasting at least 72h (**Fig. 1**). In the reversal task, the active port switched every 25 pellets collected, thus creating a new behavioral block. Performance within blocks was quantified using two FEDUPP-derived metrics (**Fig. 5)**. The first metric, ‘learn score’, measured cumulative accuracy within a block, calculated as the proportion of correct (‘active’ port) pokes from block onset up to each nose-poke event. This measure was time-invariant and reflected the progressive acquisition of the correct port over the course of a block. The second metric, ‘learn result’, calculated accuracy over the final 25% of pokes in a block, providing a snapshot of end-of-block performance once adaptation has occurred.

**Figure 5.**
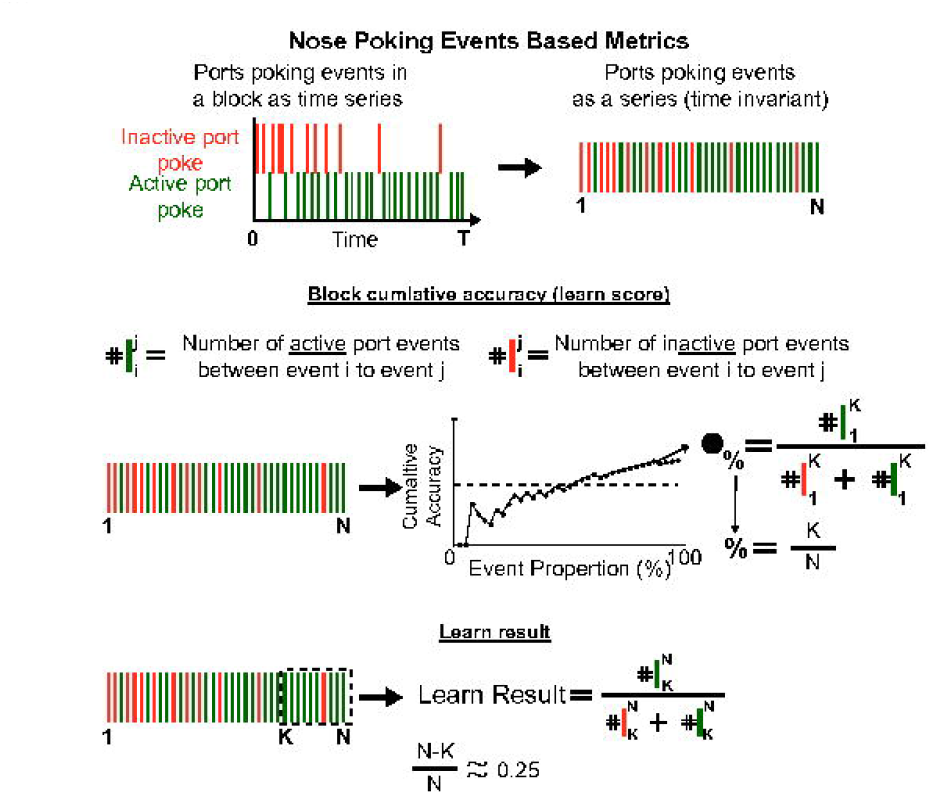
Schematic of FEDUPP calculations of nose poking events based matrix. Upper panel - Given a time series of ‘active’, ‘inactive’ port pokes (green and orange) with total N events between 0 to T, it was turned into N N-length vector, which is time invariant. Middle panel- the ‘learn score’ was the accuracy of the first K events in the N-length vector where K < N. Learn score is plotted as a function of K divided by N, which is the event proportion. Lower panel - the ‘learn result’ was the accuracy of the last N-K events in the N-length vector where K < N.

The WT cohort showed that, over the 3-day reversal phase, mice completed an average of ∼23 (23.11±0.56, N=18, **Fig. 6A**) blocks, during the 72 hours of the task, and collected an average of ∼195 pellets per day (195.42±9.04, N=18, **Fig 6B**). The learn score progressively increased the more nose pokes the mouse performed during a block (**Fig. 6C**), resulting in total accuracy for a block being just above chance level (54.35±0.60%, N=18, **Fig. 6D**). The learn result averaged ∼67% (**Fig. 6E**, 67.41±0.87%, N=18), well above the chance level.

**Figure 6.**
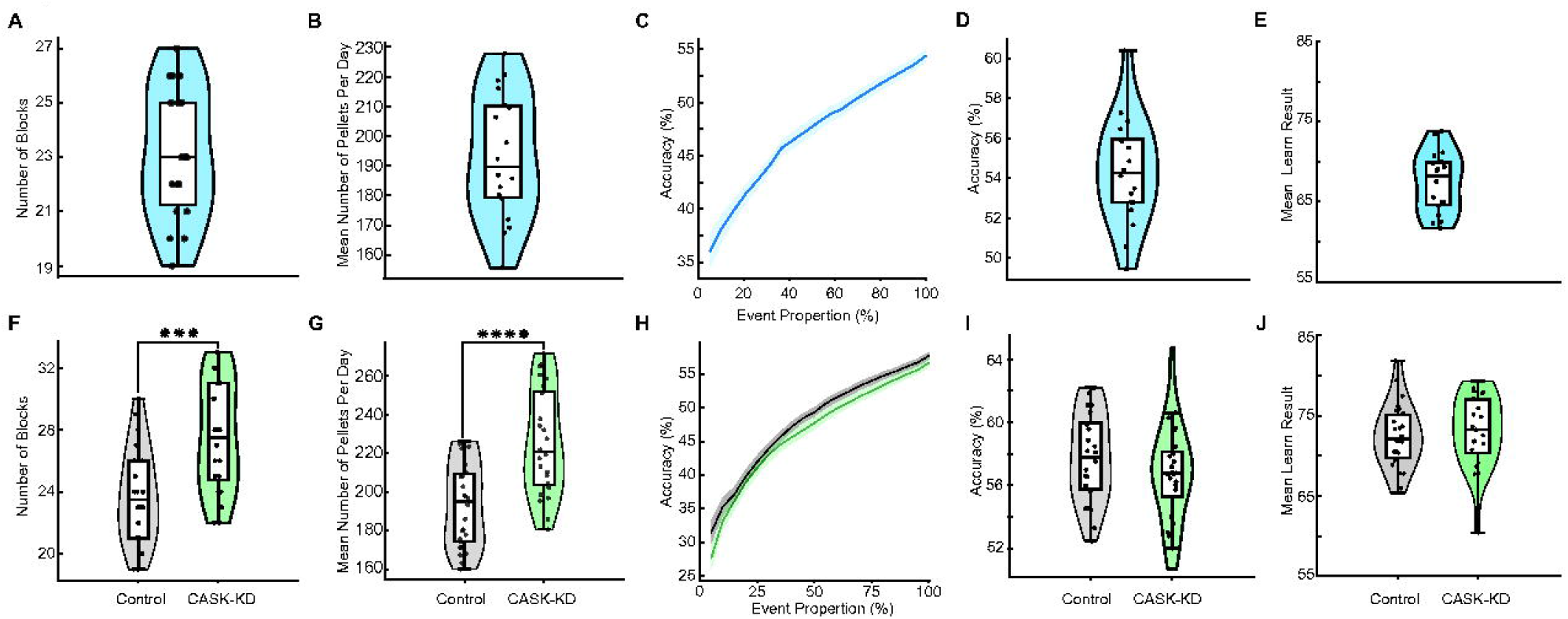
Statistics of performance during the reversal task block. **A, F**. Average number of blocks done during the reversal task for the WT (A) and Control and CASK-KD cohorts (F). **B, G**. Average number of daily collected pellets during a reversal task for the WT (B) and control and CASK cohorts (G). **C, H**. Dynamics of the learn score during a block for the WT (C) and Control and CASK-KD cohorts (H) (gray-control, cask - green). **D, I**. The average total accuracy during blocks for WT (D) and Control and CASK-KD cohorts (I). **E, J**. The average ‘Learn Result 25%’ for blocks of WT (E) and Control and CASK-KD cohorts (J). In panels F, G, I, and J, Grey and Green designate Control and CASK-KD knockdown groups, respectively.

When the same analyses were applied to the CASK-KD and control groups, CASK-KD mice completed significantly more blocks during the reversal task (23.69±0.58 vs 27.54±0.70, CASK-KD N=24, Control N=26, p=0.0001, **Fig. 6F**) and collected more pellets per day when compared to the control group (193.32±4.2 vs 225.55±5.5, CASK-KD N=24, Control N=26, p=2.83*10^{-5}, **Fig.6G**). Despite these increases, the dynamics of the learn score metric were similar between groups (**Fig. 6H**) with total accuracy for a block being just above chance level (57.77±0.55% vs 56.71±0.62%, CASK-KD N=24, Control N=26, p=0.215 **Fig. 6I**). Similarly, both groups achieved a ‘learning result of ∼72% on average (CASK-KD: 72.40±0.76%; Control: 73.03±0.90%; CASK-KD N=24, Control N=26, p = 0.601, **Fig 6H**). This analysis indicated that all groups consistently performed above chance level within each reversal block, and that the CASK-KD and control groups had similar performance, as measured by these behavioral metrics.

### Analysis of meal structure during the reversal block reveals that accurate meals appear in reversal blocks

To determine whether structured feeding patterns emerged during the reversal task and whether these patterns reflected task accuracy, meal analysis was applied as described in **Fig. 3**. Across all cohorts, a meal was detected in 78% of reversal blocks, indicating that goal-directed feeding events were preserved during reversal performance. In the WT group, the time required to reach the first accurate meal from the start of the reversal block was ∼72 minutes (72.03±4.79 minutes, N = 18, **Fig. 7A**). When this time was normalized to the duration of each block, the mean ratio was approximately ∼56% of the block length (55.6±2.6%, N=18, **Fig. 7B**). In addition, The mean accuracy within these meals was ∼85% (85.31±0.59%, N=18, **Fig 7C**). The proportion of pellets consumed as part of a meal increased as the block progressed (**Fig. 7D)**, reaching an average of ∼80% on average (79.6±1.6%, N=18, **Fig. 7E**). This indicated that most pellets were collected during meals.

**Figure 7.**
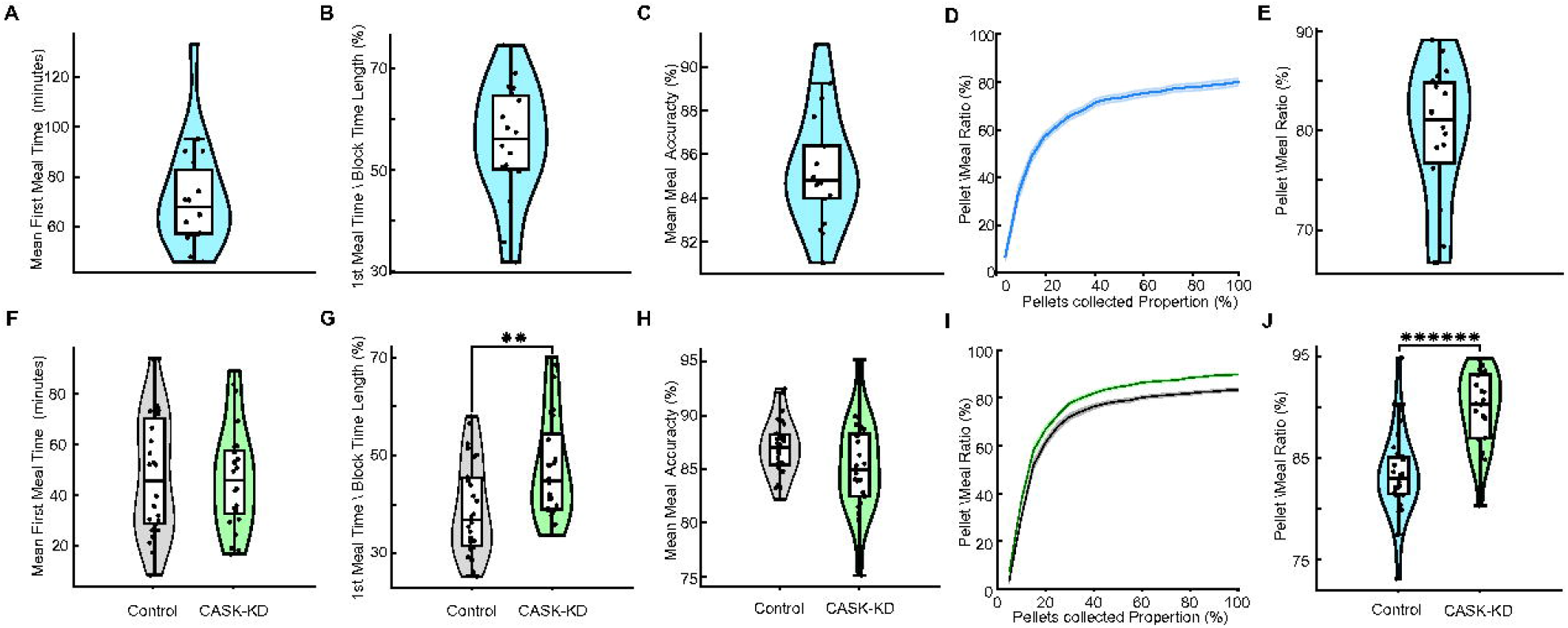
Statistics of meal properties during the reversal task behavioral block. **A, F**. Average time of first meal during a reversal task behavioral block for the WT (A) and Control and CASK-KD cohorts (F). **B, G**. Average time of first meal during a reversal task behavioral block normalized to behavioral block duration for the WT (B) and Control and CASK-KD cohorts (G). **C, H**. Average meal accuracy during the behavioral block of the WT (C) and CASK-KD (H). **D, I**. The proportion of pellets being collected as part of a meal as a function of the percentage of pellets collected during a block for WT (D) and control and CASK-KD cohorts (gray-control, cask - green) (I) **E, J**. Ratio of pellets collected as part of a meal relative to the number of all pellets collected during a block for WT (E) and Control and CASK-KD cohorts (J). In panels F-H, J, Grey and Green designate control and CASK knockdown groups, respectively.

Comparison of the CASK-KD and the control groups revealed no differences in the time required for the first accurate meals, as both groups required, on average, ∼47 minutes (CASK-KD: 46.69±4.06 minutes; Control: 47.30±4.37 minutes; p = 0.92, **Fig 7F**). However, when normalized to the length of a block, CASK-KD mice collected their first accurate meal significantly later in the block than controls (CASK-KD: 47.6±2.2%, N=24; Control: 39.1±1.8%, N=26; p=0.005, **Fig 7G**). In addition, the meal accuracy was comparable between groups (86.96±0.50, 85.27±0.84% p=, **Fig 7H**). Both groups showed an increase in the proportion of pellets consumed as part of a meal as the block progressed (**Fig. 7I**). However, the CASK-KD group reached a significantly higher average than the control with ∼90% and ∼83%, respectively (CASK-KD: 89.8±0.007%, N=24; Control: 83.4±0.008%, N=26; p=6.71*10^{-7}, **Fig. 7J**).

These findings indicate that reversal blocks are characterized by structured, accurate feeding bouts and that CASK-KD mice, while ultimately achieving similar meal accuracy, initiate accurate meals later in the block when timing is considered relative to block length, with a higher number of pellets consumed as part of the meal.

## Discussion

FED3 devices were previously used to study feeding behavior (Matikainen-Ankney et al., 2021) and, more recently, to assess cognitive flexibility either outside of the homecage sessions (Brouns et al., 2025; Murdaugh et al., 2024) or for limited home-cage sessions (Conn et al., 2024). The present study introduces two main innovations. First, a fully home-cage, two-phase paradigm was developed to continuously measure learning and behavioral flexibility by using FED3 located in the mouse homecage. In the first phase, which lasted a day, mice acquired an FR1 task, as indicated by their accuracy above chance level in selecting the active port, on average. In the second phase, the mice performed a reversal task lasting 3 days, in which the active port was switched every 25 collected pellets; thus, the reversal frequency was contingent on the mouse’s actions. Second, an open-source analysis pipeline, FEDUPP, was developed to extract multi-timescale measures of learning and cognitive flexibility from the FED3 data, including novel metrics of accuracy milestone and meal-based performance. Together, these advances allow minimally invasive, continuous tracking of operant learning and reversal behavior under naturalistic conditions.

### Estimating learning at different time scales

FEDUPP assumed that mouse FED3 nose poking represents a trial structure in which the action of a nose poke in the active port results in a pellet being released. Therefore, poking events that occurred while a pellet was available for collection were not considered to be trials when accuracy was calculated, as no pellet was released (**Fig. 1-3**). This structure creates a continuous, self-initiated trial format without explicit cues, approximating naturalistic decision-making Behavioral paradigms that measure learning and behavioral flexibility, based on consecutive trials performed by the animal, tend to run into the fundamental question “did the animal learn the task contingency?”, particularly in self-paced reversal tasks where block duration is determined by the animal’s behavior; a topic still debated (Gallistel et al., 2004; Papachristos & Gallistel, 2006; Smith et al., 2004).

To address this, FEDUPP introduced both coarse and fine temporal metrics. The overall accuracy captured performance across an entire session or block, which is a standard metric in the field (Smith et al., 2004). The ‘80% accuracy milestone’, was designed to estimate learning on a shorter time scale. This metric used a moving window of predefined time length (2h) to calculate accuracy, identifying the earliest point at which accuracy exceeds 80% within that window. This approach is conceptually similar to methods that count consecutive correct choices in a window composed of a predefined number of trials (Deliano et al., 2016; Smith et al., 2004), but with two key differences. First, FEDUPP applies a time-based window rather than a fixed trial count, accommodating a temporal variable based on the variable pace of self-initiated responding in the home cage. Second, the method applied a looser requirement for consecutive correct responses and did not conduct a formal statistical test of a null hypothesis for learning within the defined time window. This makes the metric flexible and well suited for continuous, self-paced behavioral datasets where trial structure is not externally imposed.

### Meal-based analysis as a marker of goal directed behavior

During a behavioral task, mice adopt strategies shaped by the outcomes of their actions (Griffiths et al., 2014; Hanks & Summerfield, 2017; Jackson et al., 2020). These strategies are dynamic, with individual mice shifting between exploration and exploitation during trials of a given session (Grossman et al., 2022; Le et al., 2023) based on internal states (Flavell et al., 2022) and external factors (Izquierdo et al., 2017; Woo et al., 2023). In the FR1 and reversal tasks, two features may influence a bias toward more exploration based strategies: 1) there is no cost for poking the inactive ‘port,’ and 2) the effort required to obtain a pellet (a single nose poke) is relatively low value from an ethological perspective (Salamone et al., 2018).

In this assay, higher exploratory strategy could lead to more random poking of the active and inactive ports rather than exploitation, leading to the increase of active port poking over time and apparent enhanced accuracy of mouse performance. To more precisely discern between exploration and exploitation, we focused on patterns of pellet collection that can be classified as meals, which represent motivated, increased exploitation strategy (Addicott et al., 2017). In fact, homeostatic drive increases the motivational salience of food and enhances goal-directed exploitation responding, as shown by increases in progressive-ratio breakpoints, activation of valuation circuits during feeding (Moscarello et al., 2009).

Pellets collected within 1 minute of each other were classified as part of a meal, consistent with prior FED3 literature (Matikainen-Ankney et al., 2021). In the reversal task, roughly 80% of pellets were collected within these meal bouts. We then defined “meal accuracy” as the proportion of correct nose-pokes during a meal and employed a binary classifier (see Methods) to label each meal as “accurate” or “inaccurate.” Since feeding motivation drives exploitation over exploration, meal accuracy is a sensitive proxy for task understanding. Focusing on accuracy during meals yields a more fine-grained estimate of task contingency ‘understanding’ than session-wide accuracy. For this reason, FEDUPP includes metrics such as the first ‘accurate’ meal, which can serve as a better measure of how quickly the mouse has acquired the task. Therefore, a high overall accuracy along with short time to first accurate meal suggests rapid task acquisition and consistent goal-directed responding, whereas high overall accuracy with long time to first accurate meal may indicate a slower engagement in goal-directed feeding despite correct responses elsewhere. Low meal accuracy suggests persistent exploratory behavior or weak task knowledge during motivated feeding, whereas changes in meal accuracy across blocks may reflect changes in motivation. Therefore, FEDUPP allows researchers to use meal-based metrics alongside standard accuracy measures to disentangle motivation-driven changes from true cognitive flexibility deficits.

### Effect of CASK knockdown on cognitive flexibility

CASK, a multi-domain scaffolding protein enriched at synapses, is implicated in synaptic organization, neurotransmitter release, and plasticity (Atasoy et al., 2007; Hsueh, 2006). Knockdown of CASK in the dorsal hippocampus may alter synaptic signaling required for timely incorporation of new contextual information into ongoing behavior.

In the FR1 phase, the CASK-KD mice reached the 80% accuracy milestone more quickly and achieved higher overall accuracy, suggesting increased exploitation behavior once the active port was identified. In this context, increased accuracy can be explained as reduced exploration and increased exploitation. In contrast, in the reversal task, CASK-KD mice achieved the same accuracy compared to the control group and were slower than controls when collecting the first accurate meal. In other words, the same reduced exploration that enhanced FR1 performance may have delayed recognition of the port-role change, resulting in a slower onset of accurate meals despite similar overall accuracy. This profile reflects delayed behavioral adaptation during reversal, suggesting that CASK KD in the dorsal hippocampus alters the dynamics of adapting to new contingencies without impairing the capacity to learn them, a key component of cognitive flexibility.

These findings align with previously proposed roles of the dorsal hippocampus in cognitive flexibility (Busse & Schwarting, 2016; Stuart et al., 2024), particularly in rapidly updating behavior when task rules change (den Bakker et al., 2023). Indeed, prior work suggests that enhanced instrumental performance after dorsal hippocampal lesions may arise from complex behavioral changes, such as reduced behavioral flexibility and/or altered motivation, rather than improved learning per se(Busse & Schwarting, 2016). In this context, our results support a model in which CASK contributes to hippocampal-dependent processes underlying flexible, goal-directed decision-making.

### Limitations of the paradigm and study

One of the hallmarks of learning tasks that aim to measure the cognitive abilities in mice is their reliance on a motivational state to drive a task acquisition. In the present paradigm, hunger served as the primary motivational state driver of the mouse’s engagement and performance. In the presented study, the CASK-KD mice consumed more pellets in FR1 than the controls. The current paradigm design and analysis cannot distinguish whether this difference reflects (1) increased hunger or altered metabolic state (2) changes in cognitive processes or motivation that influence task performance (Salamone et al., 2018). The behavior assay accompanying FEDUPP may integrate, in future iterations, such methods to improve interpretability of behavioral differences.

### Future directions

This study assessed learning and cognitive flexibility in mice using continuous, food acquisition-based tasks performed in their home cage, and introduced FEDUPP, an open-source software package for analyzing FED3 data. Future work could expand in two main areas. First, the behavioral assays will be expanded by varying the duration and/or the number of reversal blocks in the reversal task c to examine how task structure influences mouse performance. Second, FEDUPP will include a graphical user interface tool, enabling users to easily browse FED3 datasets, define custom metrics, such as ‘accuracy’ milestones, specify meal parameters, and choose visualizations formats. Furthermore, expansion of the analysis tools will support more advanced methods to assess learning and cognitive flexibility.

## Methods

### Mice

All experimental procedures were approved by the institutional animal care and use committee at the University of California, San Diego. Mice were housed (3–5 per cage) under a 12 h light/12 h dark cycle and provided with food and water ad libitum. Experiments were conducted using age-matched C57BL/6J mice in each cohort. 20 mice were used for method development, and 66 mice were used for the CASK-KD experiment. Due to a technical issue during the method development, 3 mice were removed from the FR1 and 2 from the reversal phases of the WT group due to unrecoverable timestamp error or overly long periods of blocking when the FED3 device dispenses a pellet. Therefore, we included 17 mice in FR1 and 18 mice in the reversal phase. For comparison analysis, in the FR1 experiment, the CASK-KD condition and control had 31 and 35 mice, respectively. In the reversal task, the CASK-KD condition and control had 26 and 24 mice, respectively. The reduced number of mice between the tasks is due to a cohort of mice that underwent only the FR1 task. In addition, due to a technical issue, one mouse in the CASK-KD had less than 2 days of data and was removed from the analysis.

### Stereotaxic surgeries

Mice were anesthetized using 5% percent isoflurane and then placed on a stereotaxic frame (Model 68803, RWD, Shenzhen, China). An incision to expose the skull was made, and the skull was leveled relative to bregma and Lambda. Bilateral Craniotomies were performed using a drill. Viral preparations (AAVDJ-CMV-LACz-U6-shCASK or AAVDJ-CMV-eGFP-U6-shCASK for the CASK-KD group, and AAVDJ-CMV-LACZ-U6-shScramble or AAVDJ-CMV-eGFP-U6-shScramble for the control group, with a titer of ∼4-5×10^12) were injected through a glass pipette using a Nanoject III (Drummond Scientific, Broomall, USA). Coordinates of injections were (AP:-2 ML:0.9 DV: −2.15, −1.65, AP:-2 ML:1.4 DV: −2.1, −1.35, AP:-2 ML:2 DV: −2.1, −1.45) with injection volume of 120-200nl in each site at a rate of 3-5 nl/sec. Following surgery, the wound was stitched and the mouse was given a subcutaneous injection of ethiqaXR (3mg/Kg). Mice were allowed to recover for at least 3 weeks before the start of the experiments.

### AAV constructs

AAV plasmid constructs were designed and cloned by VectorBuilder, and their maps and sequences can be retrieved using the vector IDs at https://vectorbuilder.com. Scramble shRNA AAV vector IDs are VB180117-1020znr and VB230831-1460avd, and shRNA-CASK AAV vector IDs are VB200205-1157jmg and VB230831-1459ard. AVV vectors included either an EGFP or LacZ reporter gene. Vectors were packaged into a virus by the Gene Transfer, Targeting and Therapeutics Viral Vector Core at the Salk Institute for Biological Studies.

### shRNA-mediated CASK-KD validation by RT-qPCR

Mouse neuroblastoma N2a cells were plated in 24-well plates and transfected with ACC plasmids encoding either shCASk or scramble shRNA control (both vectors carried an EGFP reporter) using LipoD293 (#SL100668, SignaGen Laboratories) reagent according to manufacturer’s protocol. Cells were harvested 48 hours after transfection. Total RNA was extracted using the Direct-zol RNA Kit (#R2050, Zymo Research), treated with DNAse I, and reverse transcribed using the Maxima H Minus First Strand cDNA Synthesis Kit (#K1652, ThermoFisher Scientific). qPCR was performed with SsoAdvanced Universal SYBR® Green Supermix (#1725270, Biorad) using primers specific for Cask and the housekeeping gene Gapdh, as follow: Cask-Forward = ATGGGGGTATGATTCACAGG, Cask-Reverse = CTG ATGCCATTGATTTCTCG, Gapdh-Forward = TGGCACTAGAGACGGACAGA, Gpdh-Reverse = CGTCCCGTAGACAAAATGGT. Each biological replicate (independent transfection) was run in technical replicates, and technical replicates were averaged. For normalization, Ct were converted to DeltaCt = Ct(Cask) - Ct(Gapdh). Relative expression was calculated as 2^-DDCt = mean DC(sample) - mean DCt(scramble). Data are plotted as mean +/- standard deviation of biological replicates. Statistical comparisons between shScramble and shCASK groups used an unpaired t-test.

### IHC for viral infection validation

Mice were anesthetized using IP injection of Pentobarbital (50mg/Kg) and perfused using 4% PFA. Brains were kept overnight in 4% PFA and then transferred to 30% sucrose solution until they sank. Following the brain sinking, they were coronally sliced (16 μm thickness) using a sliding vibratome (Microm HM 440E, MicroM, Germany) and kept in a cryoprotectant solution (Hoffman, Murphy and Sita, 2016). Slices containing hippocampus were mounted on glass slides and then blocked with PBST (PBS + 0.3% TritonX) + 5% NGS. Following blocking, slices were exposed to anti-GFP primary antibody (GFP 1020, Aves Labs, Davis, California, USA) at a dilution of 1:1000 overnight at 4 °C. Then, they were washed in PBST and exposed to anti-chicken 488 secondary antibody (103-545-155, Jackson Immunoresearch, Westgroove, PA, USA) for one hour at room temperature. Then, they were washed and stained with DAPI (1.2 μM, Sigma Aldrich) for 30 minutes. Following DAPI slides were washed and then coverslipped using mounting media (DAVCO, Sigma Aldrich). Slices were imaged using a Keyence BZ-X800 (Keyence, Itasca, IL, USA.).

### FR1 and reversal behavioral protocols

Operant learning was evaluated using the FED3, which allows mice to be trained to nose-poke for food pellets with minimal experimenter involvement. *FED3 devices were purchased from open-ephys (https://open-ephys.org/fed3)*. FED3 tasks were conducted with the mice housed individually, with water available ad libitum. Food was accessible exclusively through the FED3, delivering 20 mg pellets (20mg 5TUM pellets (1811143), TestDiet). Each nose-poke, in either port, and pellet retrieval event was automatically recorded by each FED3, with data saved in CSV format on an SD card. Detailed analyses of FED tasks for specific cohorts are provided below.

#### *Fixed Ratio 1* (FR1)

During the FR1 task, one of the two nose ports was designated as ‘active’; A single poke in that port resulted in a pellet being dispensed. The other port was ‘inactive’, whereas a nose poke did not result in a pellet being dispensed. Mice were engaged with the FR1 task for 24-30 hours.

#### Reversal learning

The reversal learning task was composed of consecutive blocks. During each block, the mouse performed an FR1 task, based on a specific port, either right or left, being the ‘active’ port. Once 25 pellets were dispensed, the block was switched to the next block. In the consecutive block, the mouse performed an FR1 task, but now the roles of the ports, either ‘active’ or ‘inactive’, were reversed. The reversal learning task started immediately after the FR1 task.

### Analysis programming tools

For structuring, processing, and analyzing data generated from experiments conducted with the FED3, we utilized Python (version 3.9) as the primary programming language. Key libraries included pandas, numpy, and datetime, which were employed for data manipulation and temporal handling, while matplotlib and seaborn were used for data visualization.

### Metrics used in FR1 and reversal task

We provide definitions and custom metrics developed based on both prior research and specific customizations for our behavioral dataset.

#### Meal Definition

We define a “meal” by integrating established criteria from the literature (Matikainen-Ankney et al., 2021) with conditions specific to our dataset. A meal begins when two pellet retrieval events occur within 60 seconds of each other. If a third pellet is retrieved within 60 seconds of the first retrieval, the meal is extended to include this event, forming what we call a “3-pellet meal.” This process continues as long as each new pellet retrieval occurs within 60 seconds of the initial pellet retrieval, allowing for variable meal lengths while maintaining a strict temporal boundary.

Once a retrieval event exceeds the 60-second threshold from the initial pellet retrieval, the meal extension process stops. At this point, we confirm the sequence as a meal only if the proportion of correct pokes (actions directed toward the currently active side of the apparatus) exceeds 50% of all pokes made during that sequence. This ensures that the behavior is not random and reflects genuine task engagement.

#### Reversal task behavioral block

As outlined in the experimental design, during reversal tasks, the FED3 device alternates the designated active poke after a specified number of pellet retrievals. Thus, a block encompasses all events from immediately after one active-poke switch until just before the next switch.

### First Meal Time in Each Behavioral Block

“First Meal Time in Each Block” quantifies the latency between a switch event and the initiation of the first meal in the newly established block. To calculate this, we identify the start time of the first valid meal after the behavioral block starts. Meals that began before the switch to the current behavioral block are excluded to ensure each measurement reflects new conditions post-switch. This measure is analyzed both in absolute terms (minutes since behavioral block start) and as a normalized proportion relative to the block duration.

### Accuracy measurement of a given FED3 data

For a given FED3 dataset, the accuracy is calculated as the number of nose pokes in the active port divided by the total number of nose pokes in both the active and inactive ports. Both of these events were counted when no food pallet was available to be picked up.

### 80% accuracy milestone during FR1 task

The 80% accuracy milestone marks the time at which a mouse first achieves and sustains a high level of accuracy, specifically, maintaining an accuracy rate above 80% for at least two consecutive hours.

### Learn score % during a behavioral block

The ‘learn score’ % is calculated as the accuracy in the first % nose poke actions with no pellet available (nose poke in either active or inactive port). For example, Learning Score 25% is the accuracy as measured over the first 25% of nose poke actions in a given behavioral block. This measurement is invariant to the length of the behavioral block.

### Learn result during a behavioral block

The ‘learn result’ % is calculated as the accuracy of the last 25% of nose poke actions with no pellet available (nose poke in either active or inactive port).

### Unsupervised Learning and Generalization of Meal Pattern

After calculating the accuracies for all meals in the CASK-KD group (no restriction of the first 1 or 3 days), we employ a hypothesis-driven approach to classify detected meals into “accurate” and “inaccurate” categories. To achieve this, we use meals containing 3-5 pellets from both control and experimental groups in the CASK-KD experiment to train a K-Means clustering model. The clustering process segments the data into multiple classes, and we manually identify the desired accuracy patterns discussed above as “accurate” meals, with the remaining patterns classified as “inaccurate” meals. Subsequently, we train a neural network, either CNN- or LSTM-based, to refine the classification and remove non-generalizable patterns produced by the KMeans clustering. This trained model is then applied to classify meals from experiments to evaluate its generalization performance and determine the proportion of good meals, which serves as an indicator of group learning performance.

The optimal number of clusters (k) for K-Means is determined using the Silhouette Score, a metric that evaluates how well data points are clustered together, as well as manual checking for desired clusters. Before feeding labeled meal data into the neural networks, all 1D arrays are padded with −1 at the end to standardize the array length to 4, ensuring compatibility with the model input requirements. Two neural network architectures are employed: a CNN-based model and an LSTM-based model.

The CNN-based model consists of two convolutional blocks. The first block includes a convolutional layer with 16 channels, and the second block includes a convolutional layer with 32 channels. Both layers use a kernel size of 2 and are followed by ReLU activation functions. A MaxPooling layer is applied after each convolutional block, and the final feature map is passed through a fully connected layer to map the extracted features to two output classes, representing “accurate” and “inaccurate” meals (Supp Fig. 2). The LSTM-based model comprises two LSTM layers with a hidden dimension of 400, followed by a fully connected layer that maps the output to two classes. Both models are trained using labeled meal data from the CASK experiment. Both models are trained using a batch size of 256 and an Adam optimizer (Supp Fig. 2). The learning rate of the LSTM-based model is 0.0001, and that of the CNN-based model is 0.001.

By combining unsupervised (KMeans) and supervised (CNN/LSTM) approaches, we achieved a more robust classification of meal quality, ensuring that the final model could generalize well to new data. We then used this model to refine previous analyses by examining the time to the first accurate meal in each behavioral block under reversal conditions and the first accurate meal in the FR1 session.

### Statistical Analysis

We use independent t-test for all statistical tests, and we use the SciPy implementation in Python to perform all tests.

## CRediT authorship contribution statement

**Mingyang Yao:** Conceptualization, Methodology, Data curation, Formal analysis, Software, Visualization, Writing – original draft. **Avraham Libster**: Conceptualization, Methodology, Investigation, Data curation, Formal analysis, Visualization, Writing – original draft, Writing – review and editing, Project administration. **Shane Desfor:** Investigation, Data curation. **Freiya Malhotra:** Investigation, Data curation. **Nathalia Castorena:** Investigation, Data curation. **Patricia Montilla Perez:** Investigation, Data curation. **Francesca Telese:** Conceptualization, Visualization, Formal analysis, Writing – original draft, Writing – original draft, Funding acquisition, Project administration.

## Figure Legends

**Supplementary Figure 1.**
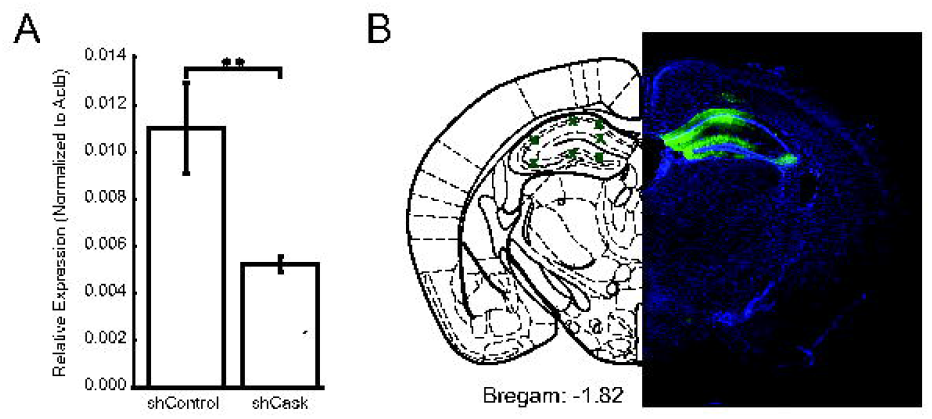
Verification of CASK knockdown. **A**. Bars are showing the relative expression of Cask mRNA levels normalized by Gapdh. Data is shown as mean +/- STDEV of 3 biological replicates.** p<0.005 unpaired t-test. **B**. Representative image of a DAPI-stained (blue) coronal slice showing the injection site into the dorsal hippocampus (green).

